# Meis1 supports leukemogenesis through stimulation of ribosomal biogenesis and Myc

**DOI:** 10.1101/2022.02.03.478956

**Authors:** Maria-Paz Garcia-Cuellar, Andreas Prinz, Robert K. Slany

**Affiliations:** Department of Genetics, Friedrich-Alexander-University Erlangen-Nürnberg, Germany

**Keywords:** acute myeloid leukemia, homeobox transcripton factor, Meis1

## Abstract

The homeobox transcription factors HoxA9 and Meis1 are causally involved in the etiology of acute myeloid leukemia. While HoxA9 immortalizes cells, cooperation with Meis1 is necessary to induce malignancy. Here, we apply degron techniques to elucidate the leukemogenic contribution of Meis1. ChIP-seq demonstrated that Meis1 localized mainly to H3K27ac and H3K4me1 modified enhancers pre-bound by HoxA9. HoxA9 was epistatic to Meis1 as degradation of HoxA9 caused an immediate release of Meis1 from chromatin. Nascent-RNA sequencing revealed the Meis1 gene expression pattern to be dominated by *Myc*, ribosome biogenesis and rRNA synthesis. While Myc accounted for cell-cycle stimulation, it could not substitute the leukemogenic effects of Meis1. Enhanced ribosomal biogenesis was accompanied by elevated resistance against RNA polymerase I and translation blocking inhibitors without affecting steady-state protein synthesis. HoxA9 and Meis1 protein stability was controlled by casein kinase 2 (CK2). CK2 inhibition caused rapid degradation of HoxA9 and Meis1 suggesting a potentially exploitable regulatory pathway.

## Introduction

Besides their function during embryogenesis, Hox-homeobox transcription factors are well established oncoproteins in acute leukemia. Particularly, HOXA9 is frequently overexpressed in hematopoietic malignancies and HOXA9 expression is an independent negative prognostic factor ^1, 2^. Reflecting the tendency of homeobox proteins to form heteromultimers, overexpression of HOXA9 in leukemia is almost always accompanied by equally elevated levels of MEIS1 and PBX3, two members of the “three amino acid loop extension” (TALE) homeobox family. Biochemical evidence showed that the PBX/MEIS interaction is required for nuclear import of the dimers ^3^. In addition, Pbx3 protects Meis1 from proteosomal degradation ^4^. As a consequence of this molecular cooperation experimental introduction of HoxA9 immortalizes hematopoietic precursor cells, but HoxA9 alone does not induce aggressive disease. Full leukemogenesis requires addition of Meis1 and can be exacerbated further by increasing Pbx3 ^4, 5, 6, 7, 8^. While known for a long time a detailed molecular explanation for this enhancer effect is still missing.

Previous attempts to clarify this phenomenon concentrated on a steady-state comparison of HoxA9 versus HoxA9/Meis1 expressing cells. This approach makes it hard to distinguish primary effects of Meis1 from subordinate events. Nevertheless, the gene for the receptor tyrosine kinase *Flt3* could be identified as a target gene of Meis1^9^. Abnormally active FLT3 signaling is clearly involved in the etiology of acute myeloid leukemia as demonstrated by the presence of activating FLT3 mutations in patients ^10^. Yet, Flt3 overexpression could be ruled out as reason for the Meis1 dependent leukemic enhancer effect. A complete genetic ablation of *Flt3* in animals did neither alter the incidence nor the kinetics of leukemia experimentally induced by HoxA9/Meis1 or by MLL fusions that induce strong transcription of *HoxA9/Meis1* as target genes ^11, 12^.

Hence, while *Flt3* is a suitable sentinel gene for Meis1 activity, other mechanisms must be responsible for the phenotypic outcome. A recent study ^13^ found increased Syk signaling in HoxA9/Meis1 transformed myeloid cells. Increased Syk activity indirectly recapitulated part of the Meis1 phenotype. Mechanistically, Syk activity was controlled by a PU.1/miRNA loop that was more active in Meis1 containing cells, however Meis1 did not control *Syk* transcription. In essence, a comprehensive study identifying direct and immediate Meis1 target genes is still missing, mainly, because suitable conditional Meis1 expression systems were not available.

Strikingly, hematopoietic development is especially sensitive towards perturbation of ribosomal biogenesis. This is best exemplified by the well characterized congenital anemias caused by inherited mutations in ribosomal proteins and also in ribosome biogenesis factors like Diamond-Blackfan-Anemia, Dyskeratosis Congenita, and Shwachman-Diamond Syndrome (for a review see ^14^). While initially characterized by a paucity of blood cells (anemia) many patients later go on to develop acute leukemia. This is elicited by secondary mutations occuring in hematopoietic precursors that are under continuous proliferative stress to supply the necessary number of blood cells. The conspicuous involvement of ribosomal proteins and ribosome biogenesis factors, including small nucleolar RNAs, in these syndromes has led to the general designation of “ribosomopathy” to summarize these pathologies. The list of genes involved in this disease etiology is growing with DDX41, a protein necessesary for ribosome production, as the most recent addition ^15^. It should be noted that there is no evidence that hematopoietic cells would contain more ribosomes or display a generally higher protein synthesis activity than other cells. Rather, the rapid supply of sufficient ribosomes in a short time frame is essential for rapid cell division that sustains the extraordinary proliferative activity of hematopoietic precursors. Here we demonstrate that Meis1 boosts HoxA9 activity through two major mechanisms. First, Meis1 amplifies a Myc program that is known to be pre-initiated by HoxA9 ^16, 17^ and second, Meis1 enhances ribosomal production as prerequisites for efficent leukemogenesis.

## Results

### Meis is an enhancer binding factor

To determine genome wide binding patterns by ChIP we generated myeloid precursor cell lines by retrovirally transducing primary hematopoietic stem and precursor cells (HSPC) with a combination of HoxA9, Meis1, and Pbx3, where one of the three proteins was individually HA-epitope tagged for each experiment (figure 1A). In addition, HSPC were also transduced with a combination of HoxA9 and a HA-tagged Meis1 fused to a mutated (F36V) FKBP. FKBP_F36V_ is a “degron” sequence allowing controlled elimination of the fusion protein by adding the PROTAC (proteolysis targeting chimera) dTAG13 that bridges FKBP_F36V_ and the endogenous E3 ubiquitin ligase cereblon, thus allowing rapid proteasomal degradation of the targeted protein ^18^.

**Figure 1:**
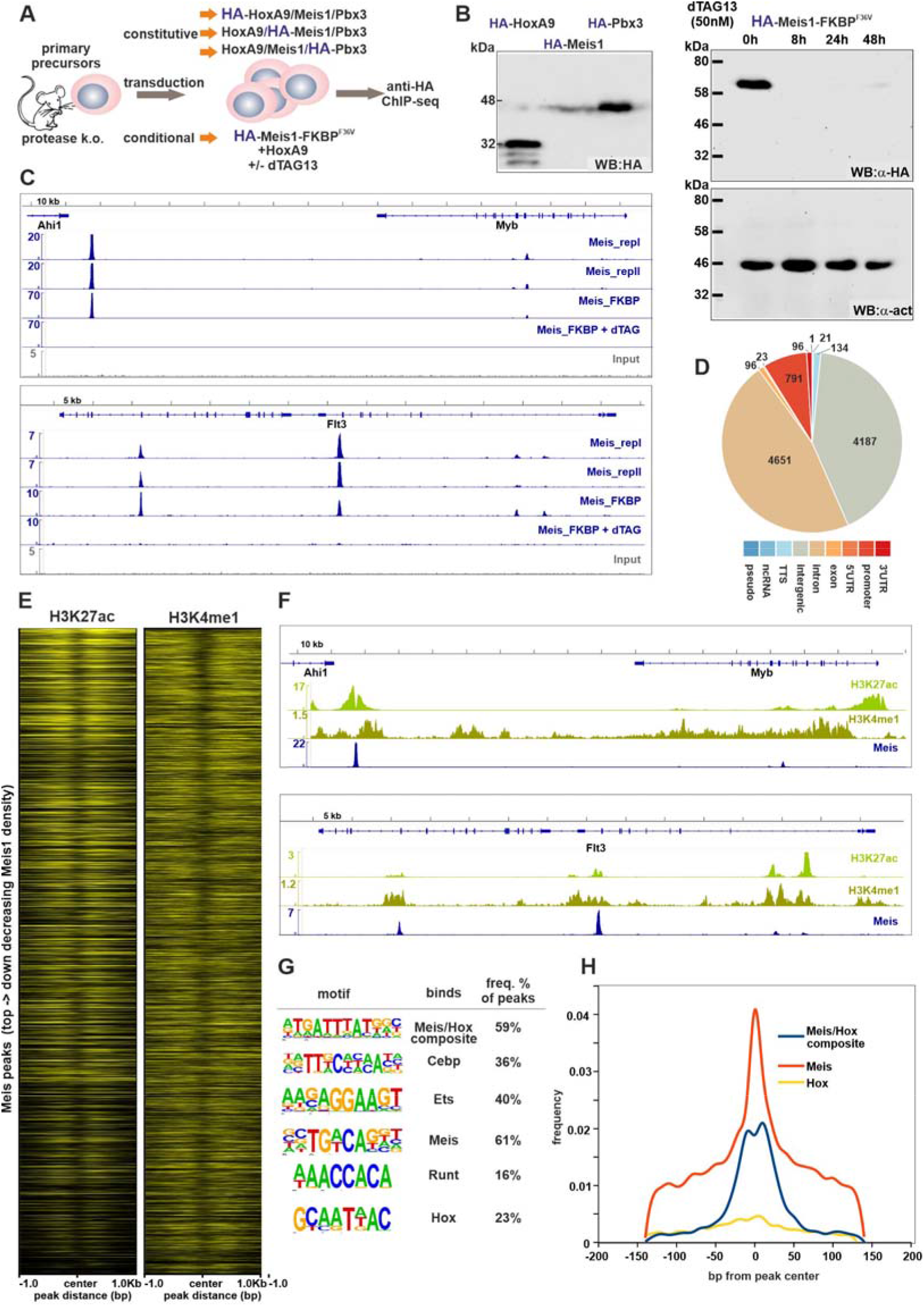
Meis1 binds preferentially to enhancers. (This figure has a supplement) A: Strategy to generate primary transformed cell lines for ChIP experiments: Hematopoietic stem and precursor cells (HSPC) were isolated from animals with a genetic knock-out of myeloid granule proteases elastase, proteinase 3, and cathepsin G to allow for efficient ChIP. B: Expression of ChIP targets and induced degradation of Meis1-FKBP_F36V_. Extracts from transformed primary cells were probed by HA-specific western for constitutive expression (left panel) of ChIP targets and for induced degradation of Meis1-FKBP_F36V_ after addition of dTAG13 (a “proteolysis targeting chimera”; PROTAC) (right panel). C: Meis1 ChIP is reproducible and peaks can be verified by Meis1 degradation. Integrated genome viewer (IGV) panels showing Meis1 binding patterns at two typical Meis1 loci, *Myb* and its major enhancer (top panel) and the known Meis1-responsive gene *Flt3* (lower panel). Tracks correspond to a replicate obtained with constitutively expressed Meis1 as well as Meis1-FKBP_F36V_ before and 8h after initiation of degradation as labeled. An input track is added as control. D: Meis1 localizes predominantly to putative enhancer locations. Pie chart of Meis1 peak distribution across functionally annotated genetic elements. E: Meis1 colocalizes with enhancer modifications. Meta-gene plots showing the distribution of enhancer-typical chromatin modifications H3K27ac (active enhancer) and H3K4me1 (putative enhancer) around the top 10000 Meis1 peaks with best reproducibility. The plot is peak centered and ordered top to down according to Meis1 binding density. F: Meis1 homes in on enhancer centers. Exemplary IGV panels demonstrating localization of Meis1 at the center of active enhancer modifications. G: Meis1 marks typical hematopoietic enhancers. *De novo* motif search results of sequences +/- 150bp of Meis1 peaks yields putative binding sites for known hematopoietic transcription factors. H: Distribution of identified binding motifs supports Meis1 or Meis1/HoxA9 composite binding at the center of identified ChIP peaks.

For ChIP HSPCs were derived from animals with a complete knock-out of the myeloid granule proteases elastase, proteinase 3 and cathepsin G (EPC mice) ^19^. This was necessary because preliminary experiments (supplemental figure 1A) showed that Meis1 is subject to rapid degradation by these proteases upon cell lysis similar to what we have described previously for HoxA9 ^17^ This proteolysis can only be stopped by rapid SDS-based denaturation of proteins but it is not inhibited by commercial protease inhibitors. Degradation also occurs during the ChIP procedure precluding efficient precipitation of Meis1-bound chromatin (supplemental figure 1 B).

Expression of individual proteins in the resulting precursor cell lines as well as functional degradation of Meis1-FKBP_F36V_ after addition of dTAG13 was checked by western blot (figure 1B) indicating that 8h after addition of dTAG13 no Meis1-FKBP_F36V_ was detectable any more. ChIP for Meis1 was performed with an anti-HA antibody in a duplicate utilizing “constitutive” cell lines, and additionally, with cells containing degradable Meis1-FKBP_F36V_ before and 8h after dTAG13 was added. Next generation sequencing (NGS) of ChIP precipitates revealed highly efficient enrichment of Meis1 bound chromatin with low background and good congruency between individual samples (figure 1C). A preliminary pass of a peak-finding algorithm resulted in more than 24000 identifiable binding sites. For further analysis we decided to concentrate on the top-scoring 10000 peaks with highest read density, because these encompassed those with the best correlation between replicates (spearman correlation 0.84 between replicates and 0.74 between Meis1 and Meis1-FKBP_F36V_, supplemental figure 1C). Reassuringly, correlation broke down (spearman correlation 0.16) after addition of dTAG13, indicating that these peaks correspond to high-confidence Meis1 binding sites.

Functional annotation revealed that the overwhelming majority of peaks (8838/10000) localized to introns or intergenic regions with only 791 binding sites annotated as “promoter” (figure 1D). As this strongly suggested a mainly enhancer-centered distribution, we determined enhancer-typical chromatin modifications H3K27ac and H3K4me1 in the same cell lines (figure 1E and 1F). As expected, these enhancer marks were highly enriched around Meis1 binding sites with Meis1 localizing to typical modification “valleys”. Generally, higher scoring Meis1 peaks corresponded also to higher H3K27ac modification levels whereas the activation independent enhancer mark H3K4me1 was present around Meis1 sites but did not correlate to Meis1 binding strength.

A *de novo* motif search of Meis1 bound sequences revealed that either Meis or Meis/Hox composite consensus binding sites were present predominantly at the center of the majority of ChIP enriched peaks (figure 1G, H). These were accompanied by consensus sites for typical hematopoietic transcription factors like Cebp, Ets- and Runt-domain containing proteins. In summary, these results point to Meis1 as a transcription factor that localizes preferentially to active enhancers in myeloid precursor cells.

### HoxA9 is epistatic to Meis1

To further elucidate the functional relationship between HoxA9, Meis1 and Pbx3 we recorded ChIP profiles for HoxA9 and Pbx3 in the newly generated Hox/Meis/Pbx lines and compared those to previously established ^17^ HoxA9 binding patterns in the absence of Meis1 (figure 2A, B). We and others have shown ^4, 7^ that Pbx3 forms physical heterodimers with Meis1 that are stable also in the absence of DNA. This was reflected in ChIP binding patterns with a nearly absolute congruency of Meis1 and Pbx3. Peaks correlated not only in location but also in binding density (figure 2C). As suggested by motif analysis, Meis1/Pbx3 peaks co-localized with areas of high HoxA9 occupancy. Despite ChIP experiments being generated in parallel by exactly the same protocol in cell lines of identical etiology, HoxA9 binding was less sharp. We noticed this phenomenon before ^17^ and this likely reflects the fact that HoxA9 has a more relaxed binding specificity, preferring AT-rich sequences. AT-rich stretches occur more frequently than the more defined Meis1 binding sites. Overall, HoxA9 binding was remarkably unaffected by the presence of Meis1. Binding profiles of HoxA9 from cells either in the absence or presence of Meis1 were superimposable and had a very good spearman correlation of 0.70 in global analysis (figure 2B). This clearly suggested that HoxA9 binding is independent of Meis1, a molecular correlate to the fact that HoxA9 alone is able to immortalize hematopoietic cells.

**Figure 2:**
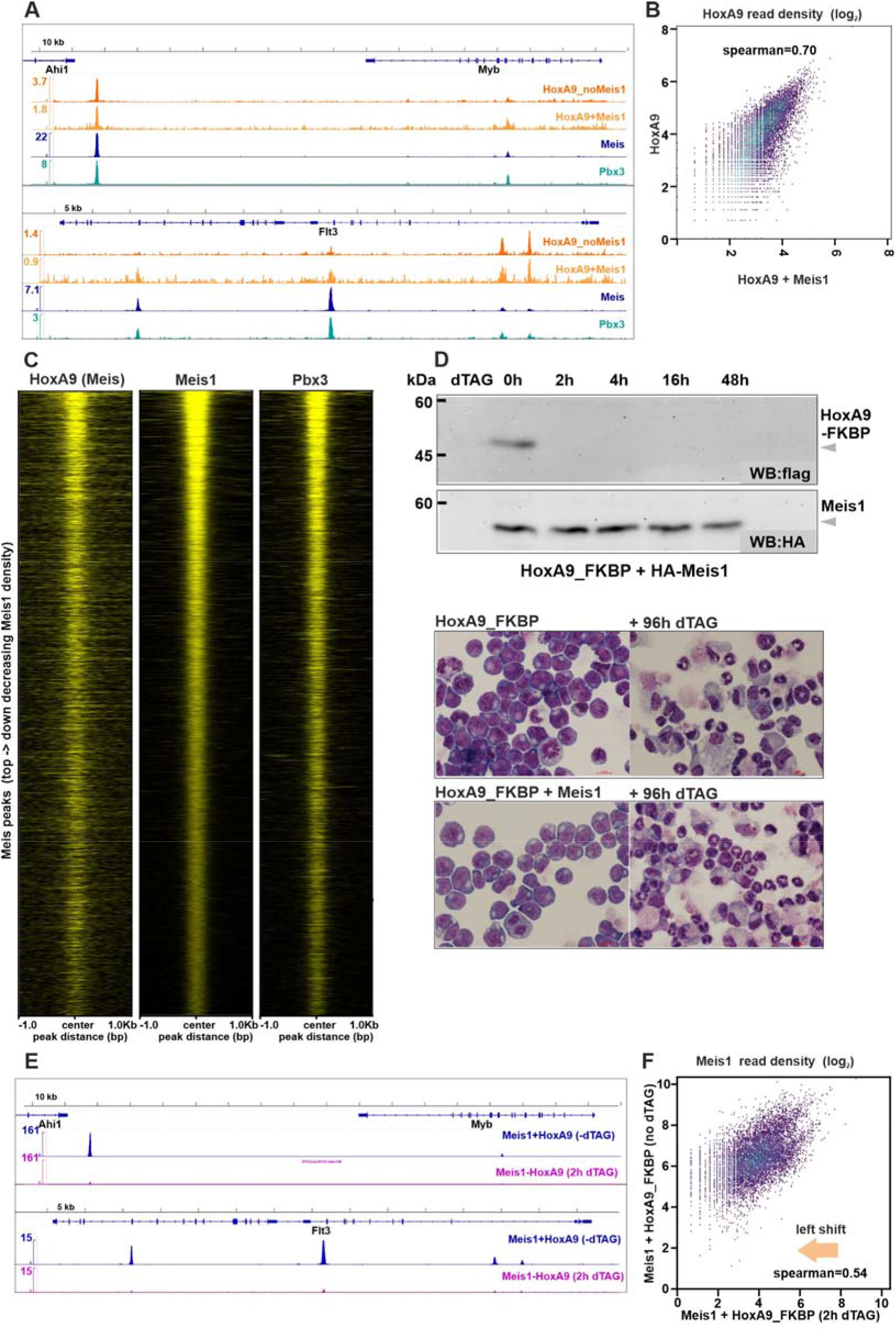
HoxA9 is epistatic to Meis1. A: Meis1 and Pbx3 bind in a more defined pattern than HoxA9. IGV plots detailing binding of Meis1, Pbx3 and HoxA9 either in cells co-expressing Meis1 (HoxA9 + Meis1) or in cells in the absence of Meis1 (HoxA9_noMeis1). Sharp colocalized peaks for Meis1 and Pbx3 are different from more diffuse HoxA9 binding characteristics. B: HoxA9 binding does not change after introduction of Meis1. Global comparison of HoxA9 binding in the vicinity of Meis1 peaks in cells transformed by HoxA9 or by HoxA9 in combination with Meis1 as indicated. C: Meis1 and Pbx3 colocalize in areas of high HoxA9 density. Metagene plots depicting binding intensity of HoxA9, Meis1, and Pbx3 around identified Meis1 peaks. Heatmaps are ordered top to down according to decreasing Meis1 binding strength. Plotted are the 10000 top-scoring Meis1 peaks as before. D: Meis1 by itself cannot maintain the transformed state of precursor cells. Upper panel: Western blot demonstrating rapid degradation of HoxA9-FKBP_F36V_ after addition of dTAG13 without affecting Meis1 protein levels. Lower panel: May-Grünwald-Giemsa stained cytospin preparations of HSPC lines generated after transduction either with HoxA9-FKBP_F36V_ alone or a combination of HoxA9-FKBP_F36V_ with Meis1. Cells are shown before and 96h after induction of HoxA9 degradation by the addition of dTAG13. Differentiation with generation of mature granulocytic cells and macrophages is observed in both cases. E: Meis1 rapidly leaves chromatin after HoxA9 degradation. IGV plots demonstrating Meis1 occupancy at *Myb* and *Flt3* loci in the presence of HoxA9 and 2h after induction of HoxA9 degradation. F: Meis1 exits from chromatin rather than changes localization after loss of HoxA9. Global Meis1 read density comparison in the presence and after degradation of HoxA9. A global left shift of Meis1 densities combined with a relative maintenance of correlation indicates loss from chromatin rather than redistribution.

To explore the reciprocal binding dependency we created a degradable HoxA9-FKBP_F36V_ fusion and used this construct to transform HSPCs alone and in combination with Meis1. Western blots (figure 2D) confirmed the rapid destruction of HoxA9 in the resulting cell lines within 2h after addition of the PROTAC while Meis1 levels were not affected. Phenotypically, degradation of HoxA9 did not cause cell death but led to a rapid differentiation of the precursor lines forming mature granulocytes and macrophages within 96h (figure 2D, lower panel). The presence of Meis1 did not alter this behavior indicating that Meis1 alone cannot maintain the undifferentiated, transformed state. At the molecular level the loss of HoxA9 was immediately followed by a loss of Meis1 from chromatin, with more than 90% of Meis1 disappearing in ChIP 2h after initiating HoxA9 degradation (figure 2E). This process did not alter Meis1 localization but rather affected the occupation density of all Meis1 peaks (figure 2F). In summary, these results are best reconcilable with a role for Meis1 and Pbx3 to “sharpen” enhancer activity on chromatin pre-occupied by HoxA9.

### The Meis1 induced genetic program is dominated by Myc and ribosomal biogenesis

Next, we wanted to identify the genes under immediate control of Meis1 by nascent RNA sequencing. This procedure (figure 3a) allows specific labeling, isolation, and sequencing of newly synthesized RNA through puls labeling of cells with 4-thiouridine reflecting thus RNA synthesis rates and transcription factor activity. Stable RNA, as determined by classical RNA sequencing, is subject to additional control mechanisms e.g. RNA export and degradation. To obtain a second independent experimental setup, we created an additional doxycycline inducible Meis1 expression system (figure 3B). Because doxycyline based retroviral expression often suffers from variegated expression, we used a fusion of a truncated LNGFR (low affinity nerve growth factor receptor) and Meis1 separated by a 2A “peptide bond breaker” (also called “self-cleavage” peptide) thus allowing for bicistronic expression. In this way, membrane-displayed LNGFR can be used to select and enrich properly expressing Meis1 cells. Western blots demonstrated the successful expression of Meis1 next to a minor amount of non-separated LNGFR-Meis1 fusion (figure 3B, lower panel). The doxycycline inducible system was transduced into wild-type HSPCs to control for any unlikely impact of the EPC protease knockout on gene expression.

**Figure 3:**
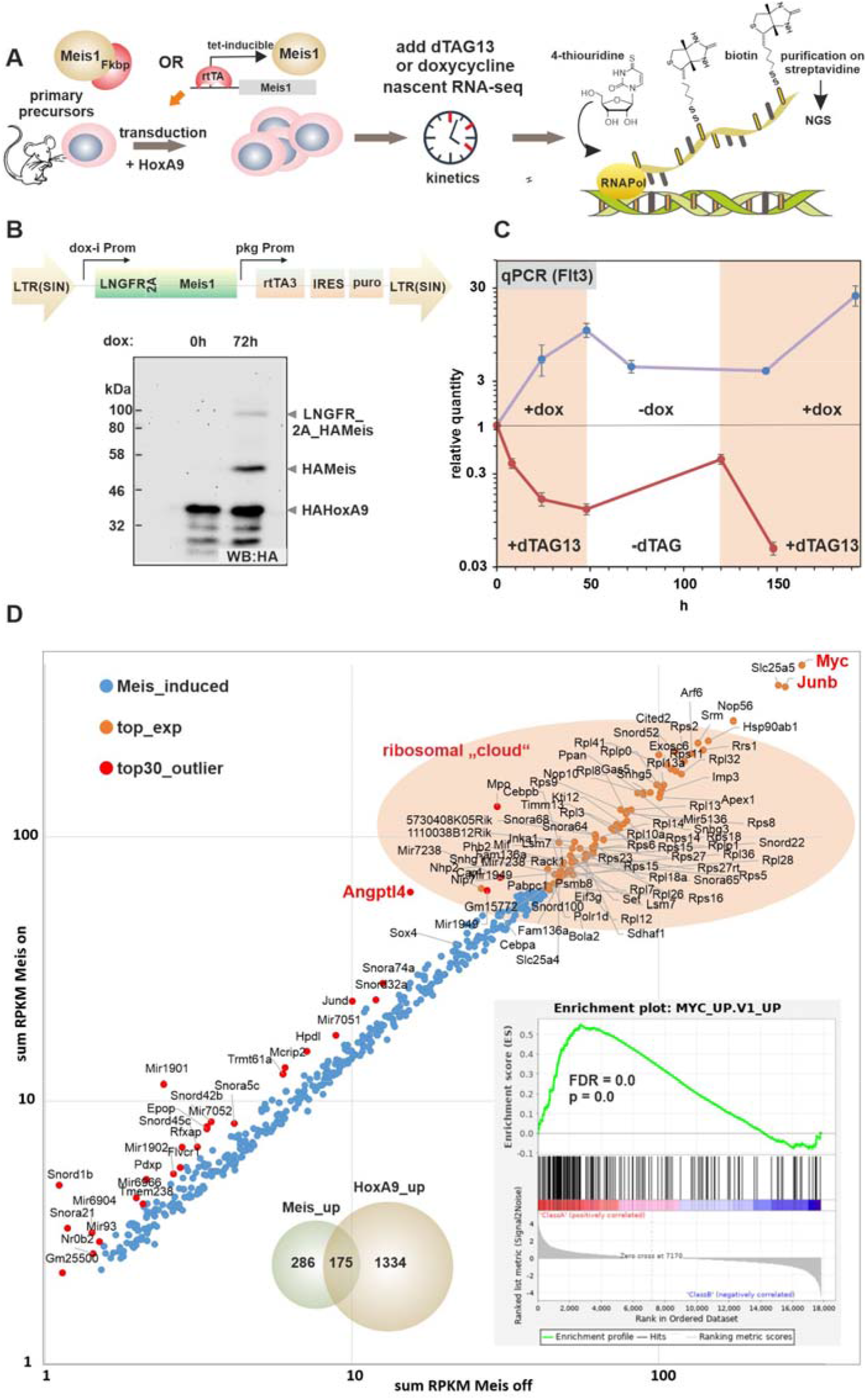
The Meis1 induced gene expression program is dominated by *Myc* and ribosome biogenesis. (This figure has a supplement and a supplemental table) A: The primary Meis1 controlled gene expression program can be determined by nascent RNA sequencing. Overview of experimental strategy. B: Meis1 expression can be positively induced in a doxycycline controlled system. Schematic depiction of an “all-in-one” inducible expression system based on a self-inactivating (SIN) retroviral backbone. LTR = long terminal repeat, self-inactivating after proviral integration, LNGFR = truncated low affinity growth factor receptor displayed on membrane for antibody-based (anti human CD271) cell selection, 2A = viral-derived “self-cleaving” peptide, blocking peptide bond formation during translation and thus allowing expression of two proteins from fusion sequence, rtTA3 = reverse tetra/doxycycline inducible transactivator of 3^rd^ generation, IRES = internal ribosomal entry site, puro = puromycine resistance. The western blot shows expression of HA-Meis before and 72h after addition of doxycycline in transduced HSPCs. Small amounts of full length LNGFR-2A-Meis1 fusion are also visible. The western was done with hot SDS extracts as the dox-inducible Meis1 system was introduced in wt cells. C: The conditional Meis1 expression constructs are biologically active. The amount of RNA coding for the Meis1 sentinel gene *Flt3* was determined by RT-qPCR in a time series either after induction of Meis1 expression by doxycyclin addition or after induction of Meis1-FKBP_F36V_ degradation by supplementation with dTAG13. To demonstrate reversibility inducers were washed out after 48h and re-added again after 120h. Values were normalized to actin transcripts and starting amounts before treatment were defined as one unit. D: Meis1 induces a Myc and ribosome-synthesis dominated gene expression program. Transcription rates in inducible Meis1 cells were determined in the Meis_on (72h dox-added or dTAG13 absent) and in the Meis_off state (dox-absent or 24h dTAG13 present) by nascent RNA sequencing. For graphical representation RPKM expression values for each gene bank accession number were added and plotted in Meis_on and Meis_off states. Shown are values (collapsed to individual genes) for all genes with a significant induction defined as log_2_(RPKM_Meis_on_/RPKM_Meis_off_)_dox_ + log_2_(RPKM_Meis_on_/RPKM_Meis_off_)_dTAG_ > 1.0. Red dots denote the top 30 outliers in gene expression change. The top 100 expressed accessions are colored orange (aggregated to gene names). Red labels identify genes investigated further. The left inset shows a Venn-diagram displaying overlap of primary Meis1 induced transcripts with HoxA9 targets identified previously ^17^ by a similar approach. The right inset depicts the top-scoring result of a gene set enrichment analysis demonstrating a strong similarity of the Meis1 induced expression pattern to the known Myc regulated program.

The dox-inducible and degron constructs were both tested for biological function by their ability to modulate transcription of the Meis1 sentinel gene *Flt3* by qRT-PCR (figure 3C). Production of Meis1 either by induction or by terminating degradation strongly induced *Flt3* whereas *Flt3* expression dropped again in the absence of Meis1.

Nascent RNA was generated before and 24h after degradation of Meis1-FKBP_F36V_ with dTAG13. Because of the considerable slower response of the dox-system, reaction time was extended to 72h for dox-addition. Transcription rates in Meis1-on and Meis1-off states were determined by next generation sequencing followed by mapping against the mm10 reference database. We filtered genes for presence (RPKM > 1) and we defined a significant change if the log_2_-fold changes between the Meis1 on/off state added up to larger than one for induced genes or smaller than minus one for repressed genes [(log2(0h/24h))_degron_ + (log_2_(72h/0h)_dox_ > 1 or < −1]. In total this yielded 492 induced genes (corresponding to 725 different GeneBank accession numbers) and 198 repressed genes (312 accession numbers) (supplemental table 1) suggesting that Meis1 acts predominantly as activator (see figure 3D for activated and supplemental figure 2A for repressed genes). Because Meis1 recognizes pre-activated enhancers rather than creating *de-novo* transcription, relative expression changes were mostly moderate. Weighing in absolute RPKM values, *Myc* was at the apex of all Meis1-induced genes. This Myc-dominance was also corroborated by GSEA (gene set enrichment analysis). The Meis1 induced gene expression program showed an unusually good match with the Myc-expression profile deposited in the molecular signature database (MSigDB) (figure 3D, inset). This fit well with strong Meis1/Pbx3 binding occuring at the known ^20^ long-range *Myc* enhancer (supplemental figure 2B). A further conspicuous feature of Meis1 controlled expression was the upregulation of a large number of genes coding for ribosomal and ribosomal biogenesis components. This included ribonucleoproteins, ribosome biogenesis regulator 1, nucleolar proteins, small nucleolar RNAs and/or their respective host genes and also subunit D of RNA Polymerase I (*Polr1d*), the sum of which we label “ribosomal cloud”. Reflecting the precursor nature of HSPCs transformed by HoxA9/Meis1, genes repressed by Meis1 encompassed differentiation promoting factors like Ngp (neutrophil granule protein), Cebpe (CAAT enhancer binding protein epsilon) and Id2 (inhibitor of DNA binding 2). In line with a physically largely overlapping binding pattern of HoxA9 and Meis1 about 40% (175/461) of all Meis1 targets, including *Myc*, have been also found to be positively controlled by HoxA9 in our previous study ^17^ (figure 3D, inset). This overlap was even greater for Meis1 repressed transcripts with 111/198 genes also showing a similar response to HoxA9 (supplemental figure 2). This supports a role of Meis1 as amplifier of a transformative state pre-established by HoxA9.

### Myc recapitulates the proliferative aspect of Meis1 expression

To elucidate the individual contribution of downstream targets to the overall Meis1 phenotype we decided to investigate three high-scoring genes for their potential impact on transformation. *Myc*, and *JunB*, were selected because they are known oncogenes and *Angptl4* codes for a soluble growth factor involved in the etiology of various solid tumors ^21^. After confirming their Meis1-dependent transcription (figure 4A), these genes were individually transduced into HoxA9 pre-transformed primary HSPCs. RT-qPCR confirmed that their expression exceeded the levels observed in HoxA9/Meis1 cells (figure 4B). We recorded proliferation, cell cycle distribution and colony forming cell (CFC) numbers of doubly transduced cells. In these experiments *Myc* fully mimicked the cell cycle and proliferation stimulation by *Meis1* (figure 4C, D) but neither *JunB* nor *Angptl4* recorded any effect. High *Myc* levels also increased CFC numbers, but due to the replicate variability in replating assays, the moderate effect of the other genes remained insignificant (figure 4E).

**Figure 4:**
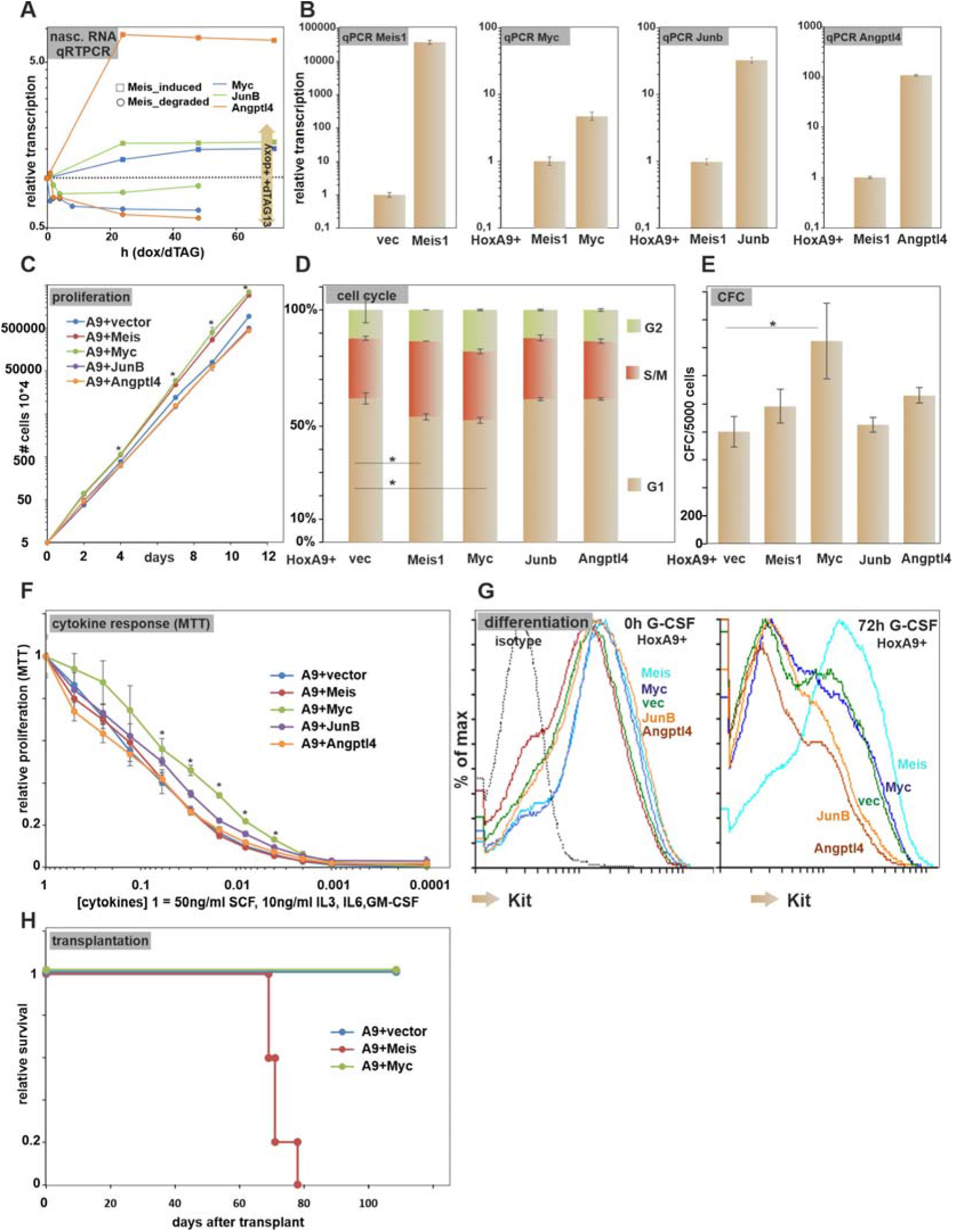
Myc controls the proliferative aspect of Meis1 activity. A: *Myc*, *JunB*, and *Angptl4* are direct targets of Meis1. Transcription rates of *Myc*, *JunB*, and *Angptl4* were determined by RT-qPCR on nascent RNA isolated in a time series after induction/degradation of Meis1. The response kinetics suggest direct control by Meis1. B: Individual expression of single target genes. RT-qPCR after transduction of HoxA9 transformed cells with individual Meis1 target genes demonstrates higher expression than in HoxA9 + Meis1 cells. C: Myc and Meis1 accelerate cell proliferation. Individual cell lines, transduced as labeled were cultivated under identical conditions and cell proliferation was determined by counting triplicates. * = p<0.05 in two-sided T-test. D: Myc and Meis1 induce cell cycle. Propidium-iodide staining of test cell lines demonstrates more cells in cell cycle (S/M and G2 phases) as consequence of Meis1 or Myc coexpression. Values correspond to averages and standard deviations of triplicate experiments. * = p<0.05 in two-sided T-test. E: Myc increases CFC numbers. Colony forming cell capacity was tested for all test lines by seeding 5000 cells in triplicate into semi solid methylcellulose medium and by counting resulting colonies after 4 to 6 days of incubation. Average and standard deviation is given. F: Meis1 does not influence cytokine signaling. Test cell lines were plated in medium supplemented with a serial dilution of cytokines (1-fold = 100ng/ml SCF plus 10ng/ml each of IL3, IL6, and GM-CSF). Proliferation/viability was tested after 72h by a standard MTT test in triplicates and plotted relative to the value in 1-fold cytokines that was set to one unit. Averages and standard deviations are plotted. Only Myc-overexpressing cells showed a significant effect (p<0.05, two sided T-test) for some values. G: Meis1 but not individual targets retard differentiation. Surface CD117/Kit expression was determined in test lines as indicated. Data were recorded in normal conditions supplemented with four cytokines and 72h after induction of forced differentiation by replacement of normal cytokine supplementation by 10ng/ml G-CSF. H: Myc does not accelerate leukemia development. Kaplan-Meier graph depicting disease free survival of sublethally irradiated syngenic animals transplanted with cell lines transduced as indicated.

Transformed cells derived from primary HSPCs need high levels of cytokines (SCF, IL-3, GM-CSF, and IL6) for viability and growth. Recording cytokine dependency in MTT viability/growth tests allows judging if *Meis1* or any of its target genes influence cellular signaling. In these experiments only *Myc* showed a tendency to alleviate dependency on cytokines. All others including *Angpltl4* coding for a known signaling molecule were inactive, thus largely ruling out an impact on signal processing as crucial for enhanced leukemogenesis.

Finally, we probed the influence of *Meis1* and the selected target genes on cellular differentiation. HoxA9 transformed cells can be forced to differentiate *in vitro* by replacing the growth cytokine mixture by G-CSF. Maturation is accompanied by a down-regulation of surface Kit (CD117) permitting to follow differentiation by FACS (figure 4G). Only Meis1 retarded differentation with surface Kit still detectable after 72h of G-CSF treatment. *Myc* did not affect differentiation while *JunB* and *Angptl4* even accelerated the maturation process. Finally, we tested if Myc accelerates leukemogensis similar to Meis1 in syngenic transplantation experiments (figure 4H). Whereas HoxA9/Meis1 transduced cells caused rapid and fully penetrant disease in the recipient animals this was clearly not the case for HoxA9/Myc cells. This result was additionally supported by a recent report of Miyamoto et al ^16^ that performed comparable experiments with longer follow-up. Thus two independent studies confirmed that Myc does not fully substitute for Meis1 in leukemogensis despite the fact that Myc can explain the proliferative aspect of Meis1 expression. Meis1 clearly has functions beyond a simple Myc-enhancer.

### Meis1 boosts ribosomal biogenesis

The salient abundance of ribosomal biogenesis components under Meis1 control led us to check if Meis1 also influences rRNA transcription. For this purpose we quantified 18s rRNA by RT-qPCR in the Meis-FKBP_F36V_ degron system during a cycle of addition/withdrawal/addition of dTAG13 (figure 5A). As ribosomal loci are not annotated in the mouse genome (version mm10) it is difficult to read out rRNA transcriptional rates from the sequencing data. Yet, qPCR confirmed that 18s rRNA concentrations clearly followed Meis1 activity. The Meis1 dependent stimulation of rRNA transcription translated to a significantly higher resistance of Meis1-containing cells against CX5461, an inhibitor of RNA PolymeraseI (figure 5B). Importantly, this was not observed with Myc-overexpressing cells thus underscoring this fact as unique property of Meis1. The presence of Meis1 also confered higher resilience to puromycine treatment (figure 5C). Puromycine is a translation chain-terminator that incapacitates actively transcribing ribosomes. Cells that are able to replace stalled ribosomes at a faster pace are expected to be more puromycine resistant. This was clearly observed for Meis1 but much less pronounced after sole expression of Myc. Notably, Meis1 did not simply induce a general tolerance against toxins because treatment of cells with the DNA-intercalating agent doxorubicine did not reveal a differential sensitivity of the respective cells (figure 5D).

**Figure 5:**
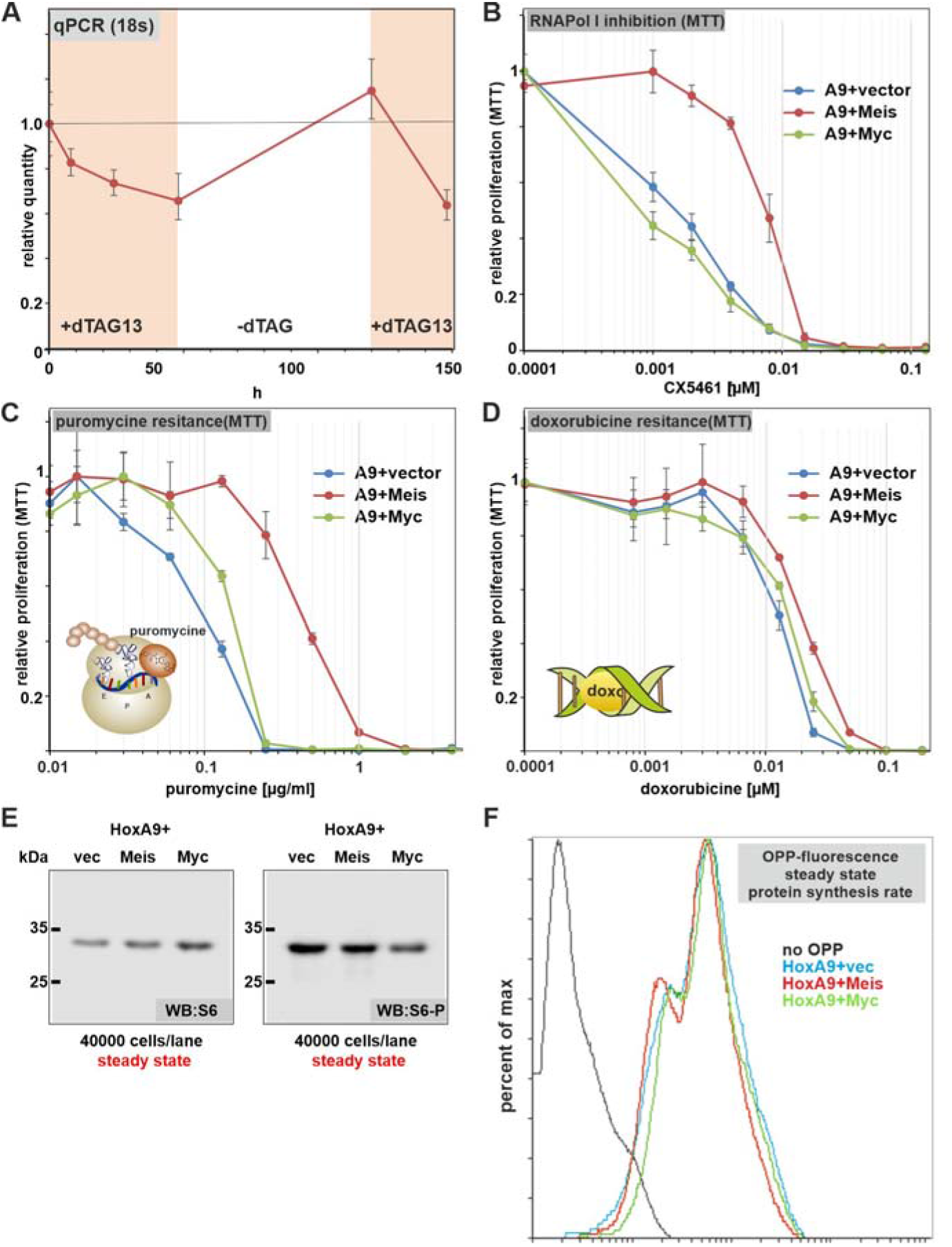
Meis1 boosts ribosomal biogenesis capacity. A: rRNA transcription correlates with Meis1 activity. Total RNA was isolated from cells transformed by HoxA9 and Meis1-FKBP_F36V_ in a time series after induction of Meis1 degradation, recovery and a second degradation phase. Concentrations of 18S rRNA were determined by RT-qPCR and are plotted relative to the starting value. B: Meis1 increases resistance against RNA Polymerase I inhibition. Cells transduced as indicated were subjected to treatment with increasing concentrations of the RNA Polymerase I inhibitor CX5461 and viability/proliferation was determined by MTT assay. Values are plotted based on untreated cells set to one unit. Averages and standard deviation of a triplicate are shown. C: Meis1 cells increases resilience towards puromycine. Experiment done as above. D: Co-expression of Meis1 or Myc does not alter sensitivity towards doxorubicine. E: Meis1 and Myc have a minor influence on steady state and phosphorylation levels of ribosomal protein S6. Cells transduced as indicated were lysed, and the equivalent of 40000 cells were loaded per lane on a SDS PAGE for detection of with S6 and phospho-S6 specific antibodies. F: Meis1 and Myc do not alter steady-state protein synthesis rate. Cells were incubated with O-propargyl-puromycine (OPP) for 30 min, fixed and then OPP was conjugated with Alexa-Fluor 488 and chain-terminated translation products were recorded by FACS.

We also checked concentration of ribosomal protein S6, its phosphorylation status, as well as overall protein synthesis rates by short-term incorporation of a fluorescently labeled puromycine derivative (OPP-puro assay). These assays did not indicate a significantly elevated number of ribosomes or an increase in overall protein synthesis capacity. A slight increase of S6 induced by Meis1 and Myc was counteracted by a concomitant reduction in phospho-S6 (figure 5E). As a consequence the overall protein synthesis activity was unchanged between the individual cell lines (figure 5F). This indicates that Meis1 enhances the synthesis rate of ribosomes rather than their final activity under steady state conditions.

### HoxA9 and Meis1 stability are regulated by phosphorylation

During the normal trajectory of hematopoietic development, cells must exit the highly proliferative precursor state and therefore they need to curb the pro-proliferative activity of HoxA9 and Meis1 at some point. Previously, we have shown that Meis1 stability is regulated by Pbx3. Pbx3 interacts with Meis1 blocking access to an ubiquitination site and as a consequence it inhibits proteasomal degradation^4^. This mechanism has a slow response rate, as it requires the cessation of Pbx3 transcription and the loss of remaining Pbx3 protein. In an attempt to identify faster acting regulatory mechanisms, we scanned the PhosphoSitePlus database (www.phosphosite.org) for known post-translational modifications of Meis1. Strong phosphorylation of Meis1 has been detected at a serine stretch between aa192 and aa 200 with the highest modification density occurring on S196 (figure 6A). Because this is immediately downstream of the Pbx3 interaction site, we decided to study the influence of phosphorylation on Meis1 stability and Pbx3 binding. For this purpose, we constructed a phosphomutant version replacing three serines against alanine (Meis_AKADA) and a phosphomimetic version (Meis_DKDD) introducing two negatively charged aspartic acid (D) residues. The effect of the mutations on stability was tested first by transfection of wt Meis1 and the respective phospho-mutants together with Pbx3 into 293T cells. A Meis1 deletion eliminating Pbx3 binding served as additional control (figure 6B). Changes of the phosphorylation site did not affect the stabilizing effect of Pbx3. Co-expression of Pbx3 increased detectable Meis1 irrespective of phosphorylation capability and was lost upon deletion of the Pbx3 interaction domain. However, phospho-site mutations had a clear independent effect on Meis1 stability. The phosphomimetic Meis-DKDD accumulated to much higher levels than wt-Meis1 whereas the phospho-defective Meis-AKADA showed reduced stability. A similar reduction of phospho-defective Meis1 amounts was also seen after transduction of HSPCs with HoxA9 and the respective Meis1 derivatives (figure 6C). In contrast to the transient system where the large amounts of protein likely cannot be fully modified, in primary cells wt-Meis1 was as stable as phosphomimetic Meis-DKDD suggesting that Meis1 is fully phosphorylated in these cells.

**Figure 6:**
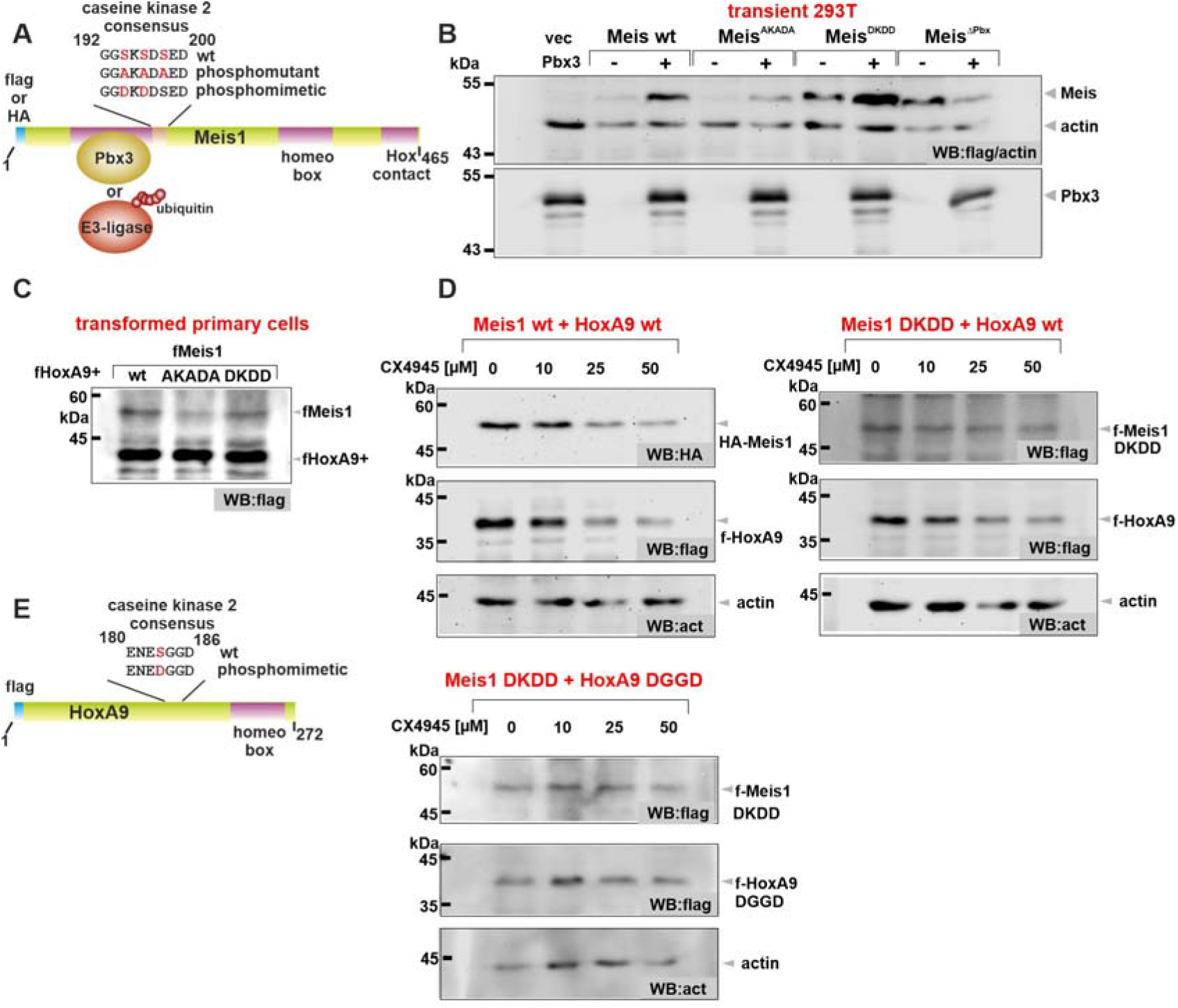
Meis1 and HoxA9 stability is controlled by caseine kinase 2 mediated phosphorylation. (This figure has a supplement) A: Meis1 contains a phosphorylation site closely downstream of the Pbx3 binding domain. Schematic depiction B: Pbx3 and phosphorylation control Meis1 stability independently. Wt-Meis1, a phosphodefective mutant (AKADA) exchanging respective serines against alanines, as well as a phosphomimetic version (DKDD) mimicking modification by introduction of negative charges were transfected with and without Pbx3 in 293T cells. Meis1 stability was recorded by western blot. Meis1 with a deletion of the Pbx3 binding domain was added as additional control. C: Meis1_AKADA is unstable in primary myeloid cells. HSPCs were transduced as indicated and the resulting cell lines were tested for Meis1 and HoxA9 expression by western blot. D: Inhibition of caseine kinase 2 reduces Meis1 and HoxA9 concentrations and phosphomimetic mutants are resistant. Primary cells transduced with wt HoxA9 and either wt-Meis1 or Meis1_DKDD were treated for 4h with increasing concentrations of the caseine kinase 2 inhibitor CX4945 as indicated. Cell extracts were probed by western for Meis1 and HoxA9. E: HoxA9 is regulated by casein kinase 2. Left panel: Schematic location of the known HoxA9 phosphorylation site. Right panel: Western made with extracts from cells transduced with phosphomimetic versions of HoxA9 (HoxA9_DGGD) and Meis1 (Meis1_DKDD) probed for HoxA9 and Meis1 after treatment with CX4945.

Next, we determined the responsible kinase for this modification. As the respective serines are embedded in a caseine kinase 2 (CK2) consensus site, we treated wt-Meis1 and phosphomimetic Meis-DKDD transduced HSPCs with increasing concentrations of a specific CK2 inhibitor (CX4945) and determined short term effects on HoxA9 and Meis1 concentrations after 4h by western blot (figure 6D). In these experiments CK2 activity correlated closely with Meis1 concentrations and introduction of the phosphomimetic changes conferred resistance to CK2 inhibition. Unexpectedly, we noted that also HoxA9 responded to a block of CK2 function. According to the PhosphoSite database HoxA9 is phosphorylated at serine 183 that is located within another CK2 consensus site (figure 6E, left panel). Alteration of this serine to aspartic acid to create a phosphomimetic HoxA9-DGGD and introduction of this mutant together with Meis1-DKDD into HSPCs yielded cells where both, HoxA9 and Meis1 were resistant to CK2 inhibition (figure 6E, right panel). Therefore we conclude that CK2 is an important natural regulator of Hox/Meis activity in primary cells. With respect to application in patients, the current general CK2 inhibitors are suboptimal. It has been estimated that CK2 phosphorylates up to 20% of all proteins in a cell ^22^. Therefore, CK2 inhibitors may have a small therapeutic window. This was reflected by the fact that cells transformed with either wt-HoxA9/wt-Meis1 or with a combination of their phosphomimetic counterparts showed an identical response towards CK2 inhibition by CX4945 (supplemental figure 3) indicating that other, unknown CK2 targets also limit overall viability after blocking CK2 in these cells.

## Discussion

Here we provide the molecular correlate of the experimental observation that Meis1 enhances leukemogenesis in combination with HoxA9 while on its own it has no discernible effect on hematopoietic development. Despite the fact that Meis1 and its binding partner Pbx3 contain autonomous homeobox DNA binding domains, HoxA9 is epistatic in hematopoietic cells. In the absence of HoxA9, Meis1 cannot maintain a stable DNA interaction and exits from chromatin. While an increased avidity for DNA of HoxA9/Meis1 dimers and HoxA9/Meis1/Pbx3 trimers was suggested previously by *in vitro* co-precipitation experiments ^4, 23^ the rapid loss of Meis1 at its binding sites *in vivo* was unexpected. Previous experiments ^24^ insinuated that dimerization of transcription factors would modify binding site specificity i.e. directing dimers to different binding sites compared to monomers. Instead, the presence of Meis1 endows enhancers pre-bound by HoxA9 with additional activity without altering HoxA9 distribution itself. This makes HoxA9 a *bona fide* pioneering transcription factor, a fact that is strongly supported by the finding that HoxA9 can induce *de novo* enhancers ^25^. HoxA9 by itself appears to have a more relaxed *in vivo* DNA binding specificity, characterized by a “diffuse” distribution. In combination with Meis1, chromatin areas are demarcated that define the center of functional enhancers. Meis1 peaks delineate the “valleys” of regions with high levels of enhancer modifications H3K27ac and H3K4me1. Thus, Meis1 focuses and concentrates enhancer activity, an effect that we call *enhancer sharpening*. It is easy to see how this process may aid to develop an increasingly active and novel enhancer architecture from previously inactive chromatin during differentiation of HSCs into highly proliferative precursors.

The effect of enhancer sharpening rather than the creation of completely new transcriptional elements fits well to the gene expression changes that we observed after induction of Meis1. Mainly, Meis1 intensified a pre-existing and primarily HoxA9 dependent gene expression pattern. Still, amplification of Myc action and the strong stimulation of ribosome biogenesis are crucial and contribute to the phenotypic manifestation of Meis1 expression. Myc, as a predominantly proliferative driver has been shown to be a major factor in the HoxA9-dependent gene expression program ^16, 17^. Cell division, however, is strongly regulated at several levels, and besides cell cycle stimulation, it requires previous cell growth i.e. an increase in cellular mass. As ribosomal proteins constitute the most abundant proteins in a cell and rRNA is responsible for > 90 percent of all RNA, synthesis of new ribosomes as preparation for actual cell division is a major bottleneck ^26^. Details about ribosome biogenesis checkpoint regulation are not yet completely clarified but it is clear that cell division cannot be executed without sufficient ribosomes. As a consequence cells with high expression of cell cycle drivers, like Myc become addicted to high ribosomal biogenesis rates, a fact that has been recently also shown for Myc induced lymphoma ^27^. The extraordinary sensitivity of hematopoietic development towards perturbation in ribosome biogenesis is underscored by the hematopoietic phenotype of ribosomopathies. Meis1 expression helps to satisfy ribosomal biosynthesis requirements of normal hematopoietic precursors and obviously also as prerequisite for efficient leukemogenesis. Strikingly, an essential step during ribosome maturation, the snoRNA guided modification of rRNA, has been recently shown to be a crucial hallmark also of the AML-ETO induced leukemogenic gene expression program ^28^.

It is still difficult to target transcription factor activity for therapeutic purposes. Elucidating how normal cells regulate transcriptional activity may reveal possible solutions to this problem. We show that posttranscriptional phosphorylation of HoxA9 and Meis1 by caseine kinase 2 is one mechanism how cells regulate activity of these two factors. This process is highly conserved during evolution as there is evidence that the insect homolog of caseine kinase 2 controls activity of the fly homeobox protein antennapedia during embryogenesis ^29^. Caseine kinase 2 is constitutively active, yet that pertains to cells in culture that are permanently cycling. It is not completely clear if this is also true for mostly non-cycling primary cells. CK2 inhibitors are tested in early clinical trials for various advanced solid and hematopoietic tumors. Results are not published yet and it will be seen if an exploitable therapeutic window exists. The finding that CK2 acts extremely pleiotropic is a caveat, but the rapid and direct response of HoxA9 and Meis1 to a small molecule treatment of an orally available inhibitor at least hints to a promising starting point for further drug development.

## Material and Methods

### DNA, cells, inhibitors, antibodies

Retroviral plasmids were constructed in pMSCV (Clontech, Palo Alto, CA) vectors. All insert sequences were either derived from laboratory stocks or amplified from cDNA isolated from murine cells and confirmed by sequencing. The FKBP_F36V_ mutation was introduced by PCR. Retroviral packaging was performed in Phoenix-E cells. HPSCs were isolated either from wt or C57/BL6 mice with a triple-ko for *Elane*, *Prtn3*, and *Ctsg* ^19^. Transduction was done with CD117 (Kit) selected cells enriched with magnetic beads (Miltenyi, Bergisch-Gladbach, Germany) essentially as recommended by the manufacturer. To generate transformed lines, cells were cultivated in methylcellulose (M3534, StemCellTechnologies, Cologne, Germany) for two rounds under antibiotics selection, then explanted and maintained in RPMI1640 (Thermo-Scientifc, Germany) supplemented with 10% FCS, penicillin-streptomycin, 5ng/ml recombinant murine IL-3, IL-6, GM-CSF, and 50ng/ml recombinant murine SCF (Miltenyi, Bergisch-Gladbach, Germany). 293T cells were from DSMZ (ACC-635 Braunschweig, Germany) and cultivated in DMEM + 10% FCS without cytokines.

CX5461 and CX4945 were purchased from SelleckChem (AbSourceDiagnostics, Munich, Germany). dTAG13 was from Tocris (NobleParkNorth, Australia). All other chemicals were provided either by Sigma (Taufkirchen, Germany) or Roth (Karlsruhe, Germany).

For western blot anti-flag M2 from Sigma (#F1804) and anti-beta actin as well as anti-HA from Cell Signaling Technologies (#3700, #3724) was applied. FACS antibody (PE-antiCD117/kit) was from ThermoScientific (eBioscience #12-1171-82). Protein synthesis rate were determined by O-propargyl-puromycine incorporation with the Click-iT^™^ Plus OPP Alexa Fluor^™^ 488 Protein Synthesis Assay Kit from Invitrogen (Waltham, MA, # C10456) according to the manufacturer’s protocol. MTT for proliferation and viability assays was purchased from Sigma and used according to standard procedures (0.33mg/ml MTT final concentration for 4h-8h cell labeling, followed by lysis in 10% SDS and readout at 550nm wave-length).

### ChIP-Seq, cell lysis, nascent-RNA isolation

ChIP was performed as described in ^30^ applying a 10 min crosslink in 1% formaldehyde @ RT followed by lysis in deoxycholate buffer (50mM Tris/HCl pH8.0, 10mM EDTA, 100mM NaCl, 1mM EGTA, 0.1% sodium-deoxycholate, 0.5% N-lauroylsarcosine 1mM PMSF and 1% HALT complete protease inhibitor cocktail (Pierce, Thermo-Fisher, Germany). Precipitation for all samples was performed with protein G coupled paramagnetic beads (Cell Signaling Technologies #9006). Antibodies used for ChIP: anti-HA rabbit monoclonal, Cell Signaling Technologies (#3724) 5μl per 5×10^6^ cells; anti-H3K27ac and anti-H3K4me rabbit monoclonals, Cell Signaling Technologies (#8173, #5326) each 5μl per 5×10^6^ cells.

Cell lysis for western was done in 20mM HEPES pH 7.5, 10mM KCl, 0.5mM EDTA, 0.1% triton-X100 and 10% glycerol supplemented with 1mM PMSF and 1% HALT complete protease inhibitor (triton lysis) or in hot (95°C) 50mM TrisHCl pH6.8, 0.2% SDS followed by a 2min nucleic acid digestion at RT with 10 units of benzonase after supplementation with 0.5mM MgCl_2_ (SDS lysis). Nascent-RNA isolation was done exactly as described in ^31^.

### NGS and bioinformatics

ChIP sequencing libraries were prepared using NEBNext^®^ Ultra^™^ II DNA Library Prep Kit reagents (NEB, Ipswitch, MA) according to the procedure recommended by the manufacturer. Size selection was done after final PCR amplification with Illumina index primers for 14 cycles. Nascent RNA was converted into Illumina compatible libraries with NEBNext^®^ Single Cell/Low Input RNA Library Prep reagents according to the standard protocol. Sequencing was done at the in house core facility on a HiSeq2500 instrument yielding 100bp single- or paired-end reads.

Data were mapped with BWA mem (0.7.17) ^32^ to the *Mus musculus* mm10 genome. Reads mapping more than once were excluded by filtering for sequences with a mapping quality score > 4. For visualization BAM files were normalized and converted to TDF format with IGV-tools of the IGV browser package ^33^. Peak finding, motif analysis and peak annotation was done with Homer (4.9.1) ^34^. BAM files were converted to bigwig by Deeptools (3.0.0, bamCoverage) ^35^. Metagene plots were created with Deeptools (3.0.0). Matrices were calculated with calculateMatrix and plotted with plotHeatmap from the Deeptools suite. RNA derived reads were aligned with STAR (v020201) ^36^ to the reference genome mm10 and reads derived from repetitive sequences were excluded by samtools (view)1.8 ^37^. Transcripts were quantified by Homer analyzeRNA routines and further analyzed with standard spreadsheet tools.

### Transplantation experiments

For transplantation experiments C57/BL6 mice were sublethally irradiated with 6Gy gamma-radiation. Syngenic cells for transplant were prepared by retroviral transduction followed by antibiotic selection during two rounds of methocel culture as above. Animals were injected 24h after irradiation intravenously with 0.5×10^6^ transduced cells and 0.5×10^6^ total bone marrow cells for radiation protection. Drinking water was supplemented with antibiotics for one week after transplant (1.1g/L Neomycin, 10^6^ units/L PolymixinB), thereafter recipients were kept with regular water and chow *ad libitum*. Daily monitoring for signs of disease (reduced activity and body hygiene, weight loss, hunched posture, altered breathing frequency) in combination with a standard institutional scoring system guided humane endpoint decisions. Post-mortem analysis confirmed leukemia development (enlarged spleen and liver, organ infiltration). All procedures were approved by the proper institutional and state authorities and license numbers are available on request.

### Data availability

Raw NGS reads were submitted to the European Nucleotide Archive under accession number ERP134562

### Statistics

Where appropriate two-tailed T-test statistics were applied.

## Acknowledgements

We thank Renate Zimmermann for technical assistance. This work was supported by research funding from Deutsche Krebshilfe grant 70114166 and in part by the Deutsche Forschungsgemeinschaft grant SL27/9-2 both awarded to RKS.

## Author contributions

MPGC, and RKS performed and analyzed experiments. AP cloned and tested degron constructs. RKS performed NGS data analysis, conceived and supervised experiments, RKS wrote the manuscript. All authors read and discussed the manuscript. MPGC, AP and RKS have no conflict of interest to declare.

## References

1. Collins CT, Hess JL. Deregulation of the HOXA9/MEIS1 axis in acute leukemia. Curr Opin Hematol 23, 354–361 (2016).

2. Collins EM, Thompson A. HOX genes in normal, engineered and malignant hematopoiesis. Int J Dev Biol 62, 847–856 (2018).

3. Berthelsen J, Kilstrup-Nielsen C, Blasi F, Mavilio F, Zappavigna V. The subcellular localization of PBX1 and EXD proteins depends on nuclear import and export signals and is modulated by association with PREP1 and HTH. Genes Dev 13, 946–953 (1999).

4. Garcia-Cuellar MP, Steger J, Fuller E, Hetzner K, Slany RK. Pbx3 and Meis1 cooperate through multiple mechanisms to support Hox-induced murine leukemia. Haematologica 100, 905–913 (2015).

5. Dickson GJ, et al. HOXA/PBX3 knockdown impairs growth and sensitizes cytogenetically normal acute myeloid leukemia cells to chemotherapy. Haematologica 98, 1216–1225 (2013).

6. Li Z, et al. PBX3 and MEIS1 Cooperate in Hematopoietic Cells to Drive Acute Myeloid Leukemias Characterized by a Core Transcriptome of the MLL-Rearranged Disease. Cancer Res 76, 619–629 (2016).

7. Li Z, et al. PBX3 is an important cofactor of HOXA9 in leukemogenesis. Blood 121, 1422–1431 (2013).

8. Thorsteinsdottir U, Kroon E, Jerome L, Blasi F, Sauvageau G. Defining roles for HOX and MEIS1 genes in induction of acute myeloid leukemia. Mol Cell Biol 21, 224–234 (2001).

9. Wang GG, Pasillas MP, Kamps MP. Meis1 programs transcription of FLT3 and cancer stem cell character, using a mechanism that requires interaction with Pbx and a novel function of the Meis1 C-terminus. Blood 106, 254–264 (2005).

10. Ambinder AJ, Levis M. Potential targeting of FLT3 acute myeloid leukemia. Haematologica 106, 671–681 (2021).

11. Morgado E, Albouhair S, Lavau C. Flt3 is dispensable to the Hoxa9/Meis1 leukemogenic cooperation. Blood 109, 4020–4022 (2007).

12. Staffas A, et al. Upregulation of Flt3 is a passive event in Hoxa9/Meis1-induced acute myeloid leukemia in mice. Oncogene 36, 1516–1524 (2017).

13. Mohr S, et al. Hoxa9 and Meis1 Cooperatively Induce Addiction to Syk Signaling by Suppressing miR-146a in Acute Myeloid Leukemia. Cancer Cell 31, 549–562 e511 (2017).

14. Sulima SO, Hofman IJF, De Keersmaecker K, Dinman JD. How Ribosomes Translate Cancer. Cancer Discov 7, 1069–1087 (2017).

15. Chlon TM, et al. Germline DDX41 mutations cause ineffective hematopoiesis and myelodysplasia. Cell Stem Cell 28, 1966–1981 e1966 (2021).

16. Miyamoto R, et al. HOXA9 promotes MYC-mediated leukemogenesis by maintaining gene expression for multiple anti-apoptotic pathways. Elife 10, (2021).

17. Zhong X, et al. HoxA9 transforms murine myeloid cells by a feedback loop driving expression of key oncogenes and cell cycle control genes. Blood Adv 2, 3137–3148 (2018).

18. Wu T, Yoon H, Xiong Y, Dixon-Clarke SE, Nowak RP, Fischer ES. Targeted protein degradation as a powerful research tool in basic biology and drug target discovery. Nat Struct Mol Biol 27, 605–614 (2020).

19. Guyot N, et al. Unopposed cathepsin G, neutrophil elastase, and proteinase 3 cause severe lung damage and emphysema. Am J Pathol 184, 2197–2210 (2014).

20. Bahr C, et al. A Myc enhancer cluster regulates normal and leukaemic haematopoietic stem cell hierarchies. Nature 553, 515–520 (2018).

21. Fernandez-Hernando C, Suarez Y. ANGPTL4: a multifunctional protein involved in metabolism and vascular homeostasis. Curr Opin Hematol 27, 206–213 (2020).

22. Spinello Z, Fregnani A, Quotti Tubi L, Trentin L, Piazza F, Manni S. Targeting Protein Kinases in Blood Cancer: Focusing on CK1alpha and CK2. Int J Mol Sci 22, (2021).

23. Ryoo HD, Marty T, Casares F, Affolter M, Mann RS. Regulation of Hox target genes by a DNA bound Homothorax/Hox/Extradenticle complex. Development 126, 5137–5148 (1999).

24. Jolma A, et al. DNA-dependent formation of transcription factor pairs alters their binding specificity. Nature 527, 384–388 (2015).

25. Sun Y, et al. HOXA9 Reprograms the Enhancer Landscape to Promote Leukemogenesis. Cancer Cell 34, 643–658 e645 (2018).

26. Lessard F, Brakier-Gingras L, Ferbeyre G. Ribosomal Proteins Control Tumor Suppressor Pathways in Response to Nucleolar Stress. Bioessays 41, e1800183 (2019).

27. Domostegui A, et al. Impaired ribosome biogenesis checkpoint activation induces p53-dependent MCL-1 degradation and MYC-driven lymphoma death. Blood 137, 3351–3364 (2021).

28. Zhou F, et al. AML1-ETO requires enhanced C/D box snoRNA/RNP formation to induce self-renewal and leukaemia. Nat Cell Biol 19, 844–855 (2017).

29. Jaffe L, Ryoo HD, Mann RS. A role for phosphorylation by casein kinase II in modulating Antennapedia activity in Drosophila. Genes Dev 11, 1327–1340 (1997).

30. Milne TA, Zhao K, Hess JL. Chromatin immunoprecipitation (ChIP) for analysis of histone modifications and chromatin-associated proteins. Methods Mol Biol 538, 409–423 (2009).

31. Garcia-Cuellar MP, Buttner C, Bartenhagen C, Dugas M, Slany RK. Leukemogenic MLL-ENL Fusions Induce Alternative Chromatin States to Drive a Functionally Dichotomous Group of Target Genes. Cell Rep 15, 310–322 (2016).

32. Li H, Durbin R. Fast and accurate short read alignment with Burrows-Wheeler transform. Bioinformatics 25, 1754–1760 (2009).

33. Robinson JT, et al. Integrative genomics viewer. Nat Biotechnol 29, 24–26 (2011).

34. Heinz S, et al. Simple combinations of lineage-determining transcription factors prime cis-regulatory elements required for macrophage and B cell identities. Mol Cell 38, 576–589 (2010).

35. Ramirez F, et al. deepTools2: a next generation web server for deep-sequencing data analysis. Nucleic Acids Res 44, W160–165 (2016).

36. Dobin A, et al. STAR: ultrafast universal RNA-seq aligner. Bioinformatics 29, 15–21 (2013).

37. Li H, et al. The Sequence Alignment/Map format and SAMtools. Bioinformatics 25, 2078–2079 (2009).

