## Supplemental material for "Meis1 supports leukemogenesis through stimulation of ribosomal biogenesis and Myc"

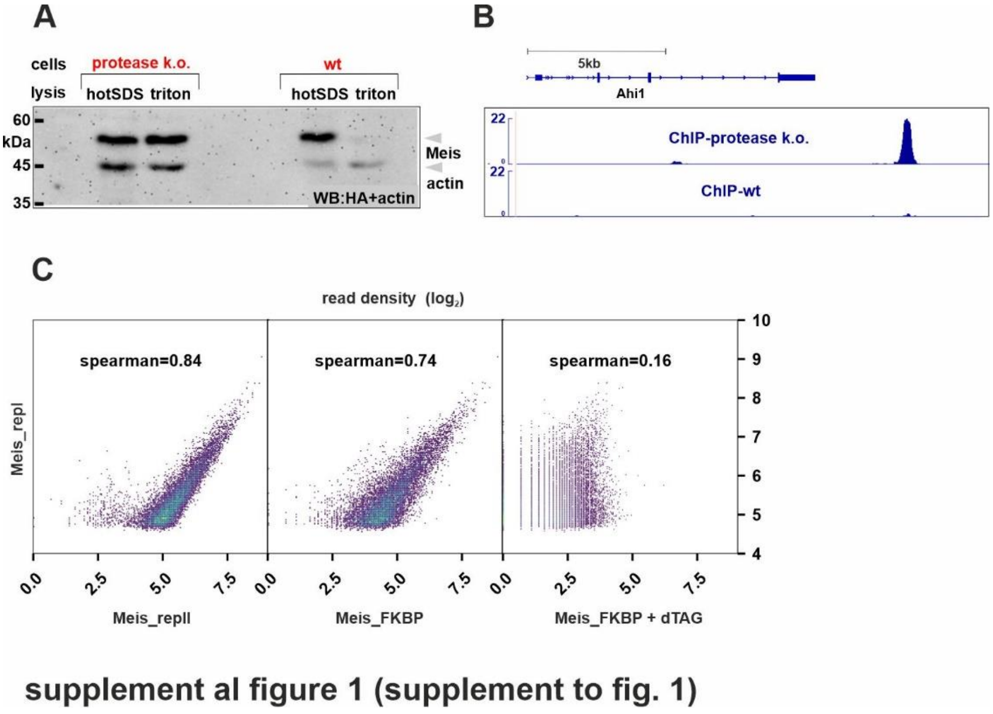


**Supplemental figure 1: Meis1 is degraded by myeloid granule proteases and can be reproducibly precipitated from protease negative cells.**

A: Meis1 is rapidly degraded in wt myeloid precursors upon cell lysis. HA (Meis1) and actin specific western blot of extracts either from wt or protease triple-k.o. HSPCs transduced with HoxA9 and HA-Meis1. Extracts were generated by lysis with hot SDS and by a triton based method (including a full complement of protease inhibitors) and probed by immunoblot for presence of HA-Meis1.

B: Meis1 can not be efficiently precipitated from wt cells. Exemplary IGV plot comparing anti-HA ChIP results generated either from wt- or granule protease knock-out cells as indicated. The exemplary Meis1 peak at the major Myb enhancer is shown.

C: Meis1 ChIP patterns are highly reproducible. Global comparison of the 10000 top-scoring Meis1 peaks comprising a defined population with high reproducibily across replicates and a Meis1-FKBP$_{F36V}$ sample. Correlation breaks down after degradation of Meis1.

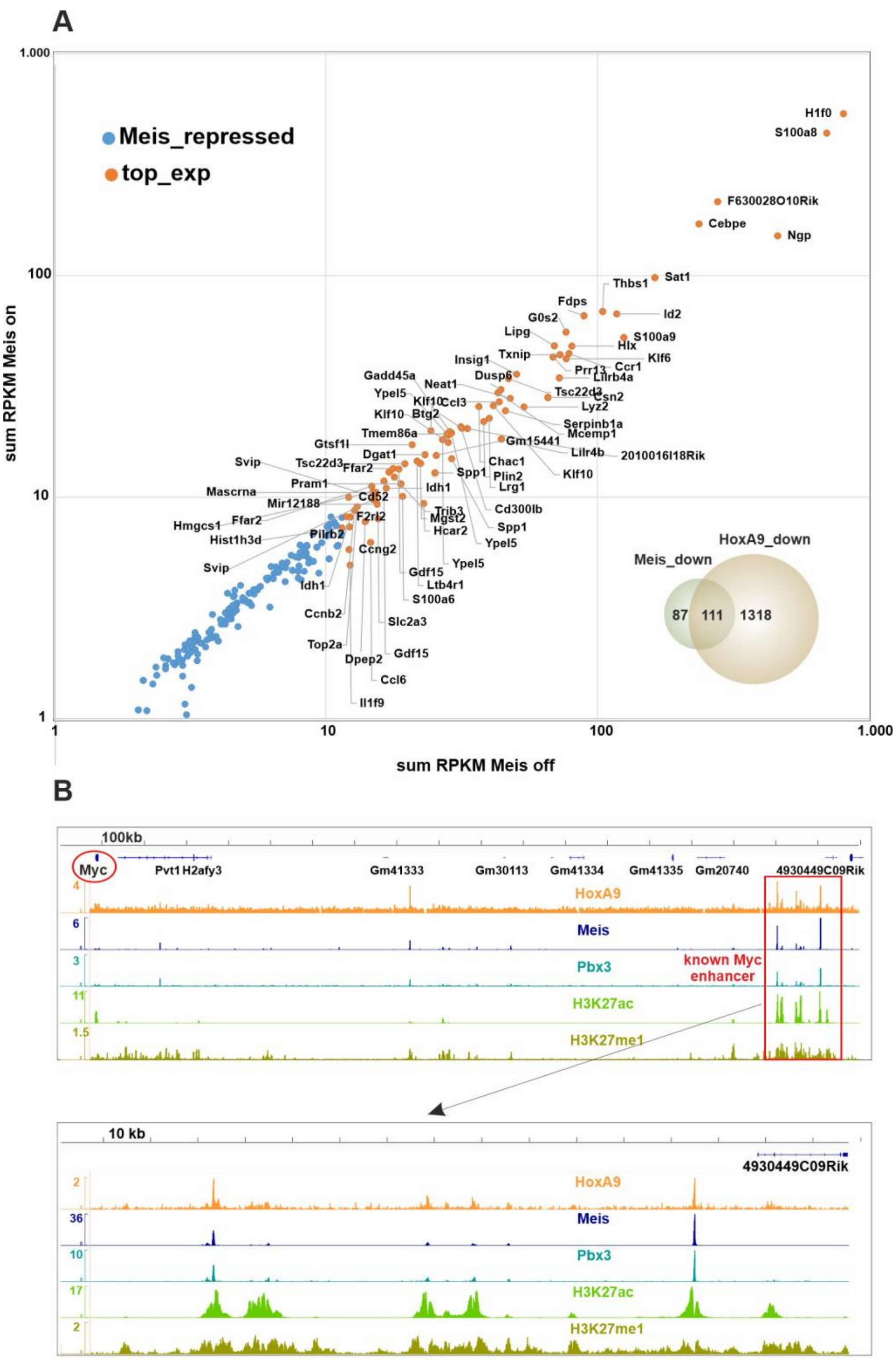


supplemental figure 2 (supplement to fig. 3)

**Supplemental figure 2: Genes downregulated after Meis1 induction, Meis1 strongly binds the known Myc enhancer**

A: Genes under negative control of Meis1. Dot plot of transcripts with reduced expression after induction of Meis1 expression. Transcript rates were determined as described for main figure 3D. For plotting and labeling individual transcripts (accession numbers) were collapsed to genes. Orange dots denote the top 100 transcripts with highest expression. The inset shows a Venn diagram detailing numbers of genes whose transcript rates were reduced by Meis1 and HoxA9 as determined previously.

B: Meis1 and Pbx3 bind to the known Myc enhancer.

Top panel: IGV plot showing the genetic environment of Myc and its major enhancer a 2MB distance that has been shown to be active in hematopoietic cells.

Lower panel: Zoom in on enhacer regions. Tracks correspond to binding of Meis1, HoxA9, and Pbx3, as well as enhancer modifications H3K27ac and H3K27me1 as indicated.

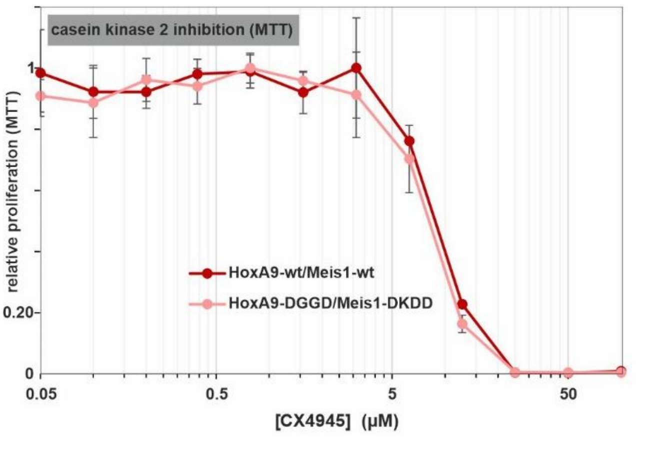


**supplemental figure 3 (supplement to fig.6)**

**Supplemental figure 3: Introduction of phosphomimetic HoxA9 and Meis1 mutants does not increase overall resistance of cells against caseine kinase 2 inhibtion.**

Cells transduced either with wt-versions or with phosphomimetic variants of HoxA9 and Meis1 as indicated were subjected to varying concentrations of the caseine kinase inhibitor CX4945 for 72h and viability/proliferation was tested by MTT assay. Relative values are plotted with untreated cells set to one unit.
